# Immune Signatures of Gastrointestinal Helminth Infection in Transcriptomes of European Bison

**DOI:** 10.64898/2025.12.04.692322

**Authors:** Mateusz Konczal, Marta Kołodziej-Sobocińska, Rafał Kowalczyk, Jacek Radwan

## Abstract

Understanding the genes and molecular pathways that shape host responses to infection is essential for advancing our knowledge of host–parasite interactions and their ecological and evolutionary implications. Such insights are especially valuable for conserving endangered species that may be vulnerable to emerging or novel pathogens. Here, we investigated the impact of the recently introduced gastrointestinal nematode *Ashworthius sidemi* on gene expression in its novel host, the European bison (*Bison bonasus*). We analyzed abomasal transcriptomes (n = 45) from individuals characterized by a wide range of infection intensities (0–8,620 parasites per host). Despite substantial variation in parasite burden, differential expression analyses detected no individual genes significantly associated with infection intensity. However, gene set enrichment analyses based on p-value distributions revealed multiple immune-related gene ontology categories, including B and T cell activation, neutrophil chemotaxis, inflammatory responses, and regulation of IL-6 production. These findings indicate that European bison mount a clear yet subtle transcriptional response to *A. sidemi* infection and highlight molecular pathways potentially involved in mediating host defense against this emerging parasite.

## Introduction

Parasitic infections can exert profound effects on host physiology, survival, and overall performance causing substantial losses in both farmed and wild populations worldwide (Coulson et al. 2018; Kaminsky and Mäser 2025). Such impacts often arise from dysregulated host immune response—either excessive, leading to immunopathology, or insufficient, resulting in uncontrolled parasite proliferation and heightened virulence. Understanding such responses is critical when a novel parasites emerge and spread across host populations that has evolved without prior exposure to such organisms, as their impact can be especially severe (Anderson and May 1986; Daszak et al. 2000). The likelihood of such events has increased in recent decades due to global changes and human activities resulting in incidental species translocations and introductions of non-native species (Patz et al. 2000; Cable et al. 2017). Vulnerability is further amplified in hosts with small effective population sizes and reduced genetic diversity, where emerging parasites can jeopardize species persistence and elevate extinction risk. These trends underscore the urgent need for evidence-based insights into host immune responses to emerging infectious diseases in endangered species, to inform conservation strategies and safeguard biodiversity (Smith et al. 2006).

Here, we investigate immunological response of European bison (*Bison bonasus L*.), an iconic species in European conservation efforts, that has recently faced the emergence of a gastrointestinal parasite, *Ashworthius sidemi*. This gastrointestinal blood-sucking helminth is primarily associated with several Asiatic cervids, including sika deer. Around 150 years ago, sika deer were introduced to many

European countries, likely bringing their parasites with them (Dróżdż et al. 1998). Even tough introduced sika deer lost their original parasites (Moskwa et al. 2014), since their introduction *A. sidemi* has been detected in several native species and has been spreading rapidly in European Cervidae populations (Magdálek et al. 2022). Unexpectedly, ashworthiosis has also become increasingly widespread in European bison. This new host species is characterized by very small effective population size as it has been restored from few captive survivors, living around 100 years ago (Tokarska et al. 2009). However, current census population size is relatively large, including around ten thousands individuals roaming in the wild and multiple seminatural and captive populations (Kowalczyk and Plumb 2022). The first cases of *A. sidemi* were recorded in 1997 in the Bieszczady Mountains population, followed by detections in the Białowieża Primeval Forest and Knyszyn Forest (Dróżdż et al. 1998, 2003; Kołodziej-Sobocińska et al. 2023). In Białowieża Primeval Forest, ashworthiosis reached 100% prevalence within just four years, with parasite abundance peaking 6–10 years after its arrival. In nearby Knyszyn Forest prevalence soon exceeded 90% and remained relatively stable over time (Kołodziej-Sobocińska et al. 2016a, 2023). Currently, *A. sidemi* prevalence exceeds 90% in all studied European bison populations, though infection intensity differs between populations, likely due to differences in supplementary feeding and consequent feeding behaviour affecting transmissibility of the parasite (Radwan et al. 2010; Kołodziej-Sobocińska et al. 2023).

The health consequences of ashworthiosis in European bison remain less well understood. Histopathological studies have documented inflammatory cell infiltration in the abomasal walls of varying severity, along with mucosal lesions and proliferation of lymphatic follicles (Osińska et al. 2010). Kołodziej-Sobocińska et al. (2016b) showed that increasing *A. sidemi* infection intensity was associated with significant declines in blood cell counts, hemoglobin levels, and hematocrit values. These findings suggest that this nematode may substantially impair bison fitness. A key factor influencing individual outcomes is likely the effectiveness of the host immune response. To evaluate these possibilities and characterize the molecular basis of host responses, we sequenced and analyzed transcriptomes from abomasal tissues of bison with quantified infection intensities.

## Materials and Methods

### Sample collection

European bison samples were collected from natural population of Knyszyn Forest previously studied in the context of *A. sidemi* infections (Kołodziej-Sobocińska et al. 2023). Analyzed here samples were collected between 2014 and 2019. During this period European bison population increased in Knyszyn Forest from 123 individuals in 2014 to 214 in 2019 (Raczyński 2015, 2020). Studied here European bison were culled during winter seasons and parasitological examination showed that they were characterized by various infection intensity with 0 to 8620 *A. sidemi* per individual (mean = 2751, median = 2480) (Kołodziej-Sobocińska et al. 2023). The samples along with their corresponding age, infection intensity, sex and sampling year are provided in Supplementary Table 1.

Tissue samples were taken from abomasa during standard veterinary necropsies of culled animals and placed in RNAlater (Sigma). The samples were then transferred to laboratory and stored in -80°C before RNA extraction. Total RNA was extracted with RNAzol and RNA concentration, quality and integrity was assessed using Qubit Fluomenter (Thermo Fisher Scientific) and 4200 TapeStation System (Agilent Technologies). Based on RNA Integrity Number (RIN) and shape of electrophoregrams 45 samples (representing 3 uninfected and 42 infected individuals) were used for transcriptome sequencing. Sequencing of mRNA was conducted by Novogen (Munich, Germany), using mRNA library preparation with ployA enrichment and sequencing on Illumina X Plus machine (PE 150).

### Differential gene expression

Raw sequencing reads were trimmed with fastp 0.24.0 (Chen 2023) and used for quantification of gene expression using Salmon 1.10.0 (Patro et al. 2017) and annotation of the of American bison (*Bison bison*) reference genome (Genome assembly: Bison_UMD1.0). A transcript to gene table was downloaded using Biomart. Salmon was then used to quantify read counts per transcript for each sample separately and the tximport package (Soneson et al. 2016) was used to import these values into R workflow. Gene-level count data were filtered using the filterByExpr function in edgeR, to remove genes characterized by low expression (Chen et al. 2025).

Normalization factors were calculated using calcNromFactors function from the edgeR, which employs TMM method (Robinson and Oshlack 2010). Normalized log-transformed counts per million (CPM) were calculated using the cpm function from edgeR package. Principal component analysis (PCA) and hierarchical clustering were performed on log-transformed CPM values to detect outlier samples, which were removed prior to differential expression analysis.

To identify differentially expressed genes we used generalized linear model with infection intensity (number of worms), age (both continuous) and sex as predictors. Dispersion estimates were calculated using the estimateDisp function, and models were fitted using the quasi-likelihood framework in edgeR. Differential expression testing was performed with glmQLFTest. Differentially expressed genes (DEGs) were identified by comparing expression across levels of infection intensity, sex, and age. P-values were adjusted for multiple testing using the Benjamini–Hochberg method, and genes with an adjusted FDR < 0.1 were considered significant.

### Gene Ontology annotation and set enrichment analysis

Gene to GO annotation were obtained from Ensembl and GO terms with gene annotations were filtered out. Gene sets were constructed by grouping Ensembl gene IDs according to their associated GO terms, resulted in a list structure. To identify biological processes associated with differential gene expression, we applied the fgsea package (1.32.4) to performed set enrichment analysis. Genes were ranked by their negative log-transformed p-values, and fgsea was run with the minimal and maximal gene set to 10 and 500 respectively. Enrichment scores (NES) and adjusted p-values (Benjamini-Hochberg method) were computed for each GO term. The sets enriched at the top of the ranked list (with low p-values) were selected. To identify the genes within each GO term that contributed most to the enrichment signal, we extracted the leading-edge genes and examined their molecular functions in relation to their up- or downregulation.

## Results

The sequencing of 45 samples resulted in average 40.1 mln read pairs per sample (from 4.0 to 129.2 million). After adapter trimming and removing low quality reads, total number of reads were reduced by approximately 0.5% (from 3.9 to 128.4 million reads per sample). Initially, 20,862 genes were assessed and after filtering out genes with low expression, the final dataset included 13,785 genes.

Clustering of samples based on their expression profiles revealed six samples clustering apart from all other samples (Supplementary Figure 1). These samples did not share exceptional infection intensities, neither they show unique characteristics according to age, sampling date or any other feature we are aware of. Thus, we decided to remove them from the further analyses. Remaining 39 samples, do not show such strong clustering and they do not cluster according to infection status (Figure 1), suggesting that transcriptomic response to infection is subtle. Indeed, differential gene expression analysis revealed no genes differentially expressed according to infection intensity (FDR < 0.1), with the lowest adjusted FDR values above 0.4. However, we were able to identify 11 genes differentially expressed between males and females at the FDR<0.1 threshold (Supplementary Table 2) and 9 genes differentially expressed according to age of an individual (Supplementary Table 3), demonstrating power of the analyses.

**Figure 1.**
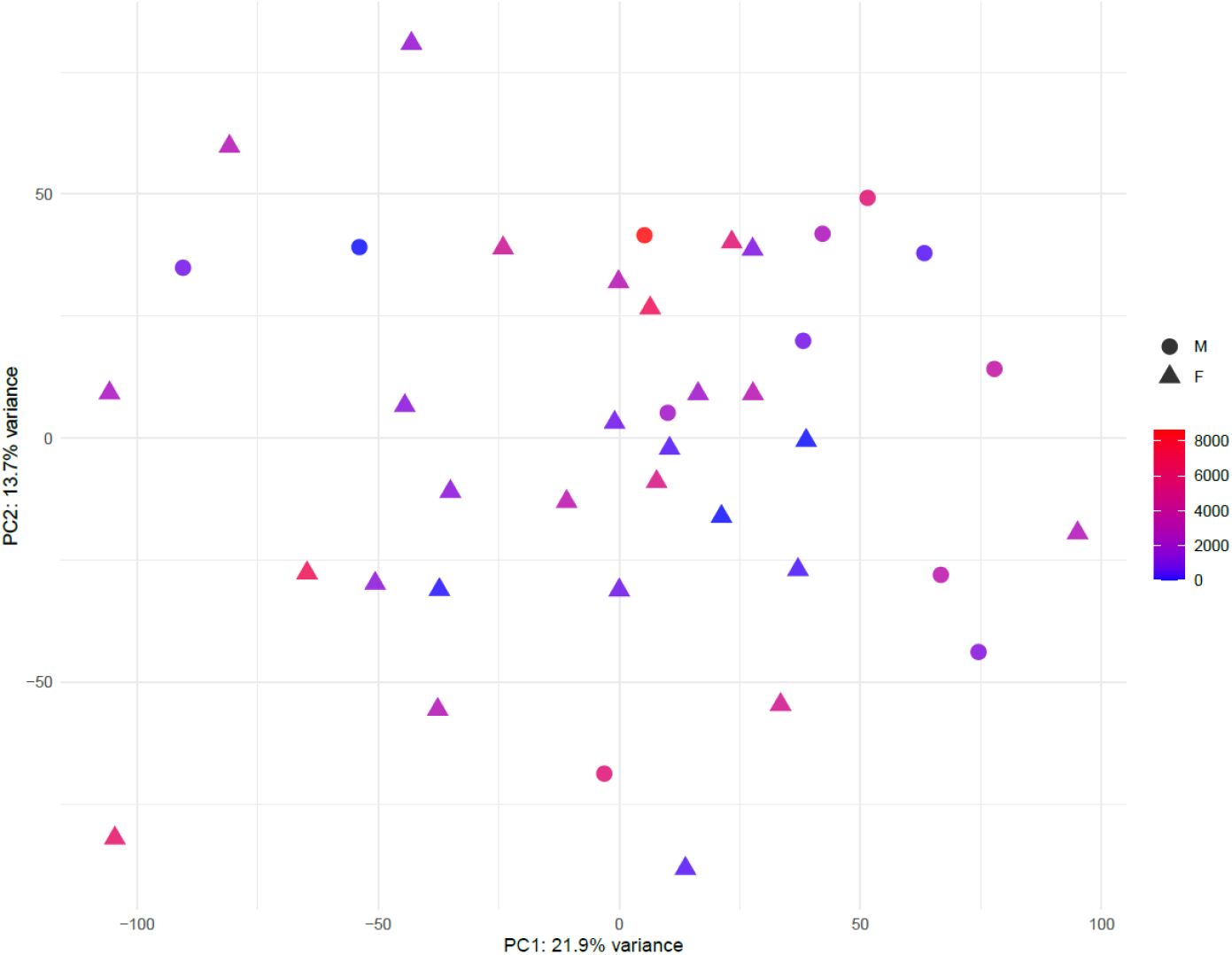
PCA of transcriptomic samples, based on their expression profiles. Males are shown as circles and females as triangles. Colors represent infection intensity measured as number of worms per individual.

Evidence that European bison in fact respond immunologically to *A. sidemi* infection was demonstrated by Gene Onotology enrichment analyses. Multiple immune-related terms were significantly enriched among genes showing low p-values according to the infection intensity, including B-cell and T-cell receptor signaling, differentiation and activation, inflammatory response, neutrophil chemotaxis or NK cell activation (Table 1). Among the leading-edge genes associated with these GO terms, those showing the largest changes in expression were mostly upregulated in highly infected individuals (Figure 2). However, when considering all leading-edge genes, the overall pattern of regulation was mixed (Supplementary Figure 5). Function of these genes and patterns of expression changes suggest active mucosal adaptive response to infection. There were no significant GO terms associated with age, and significant GO terms associated with sex represent just a couple of categories, like proteosome complex, sequence-specific DNA binding and G protein-coupled receptor binding (Supplementary Table 4).

**Table 1.**
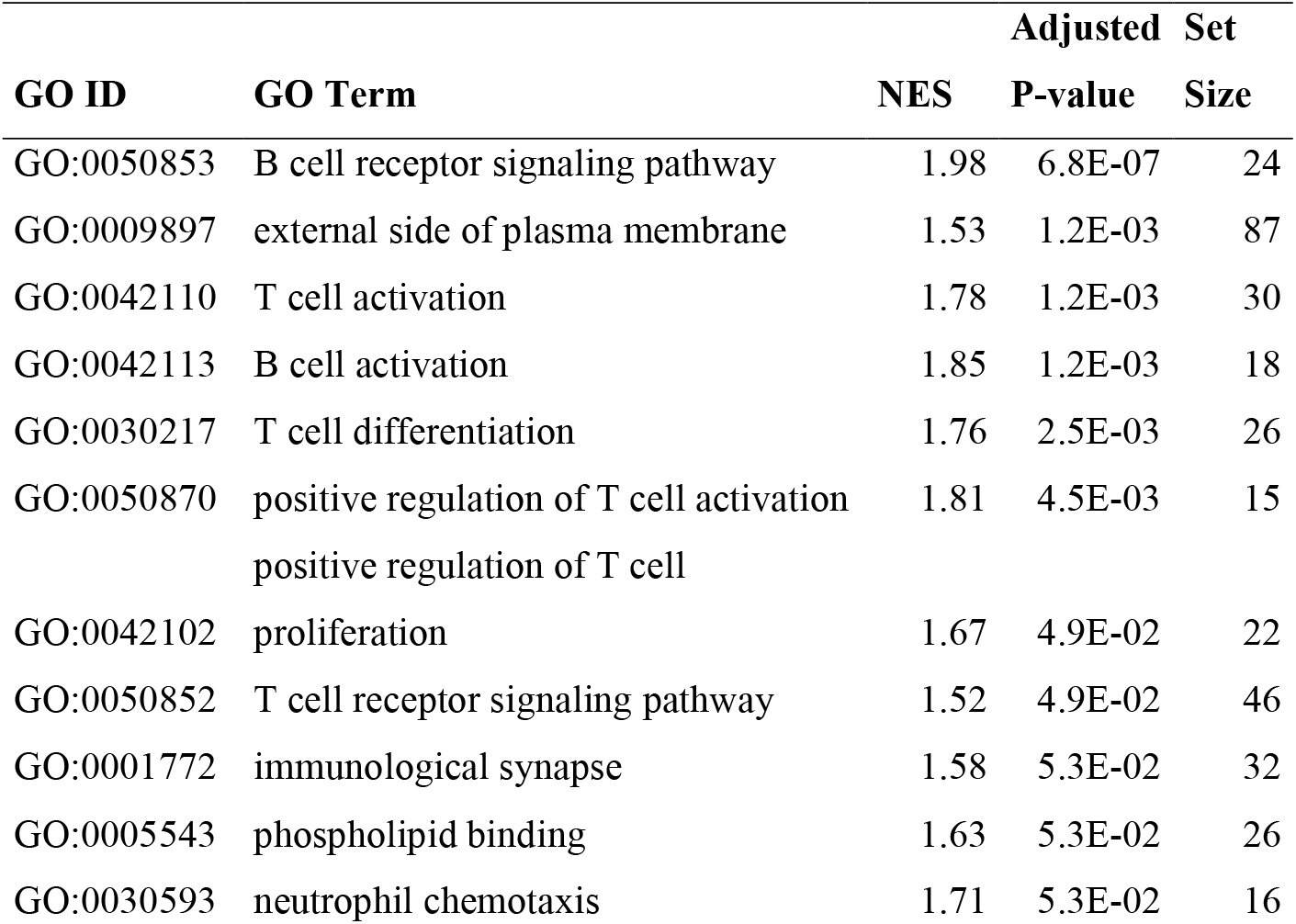
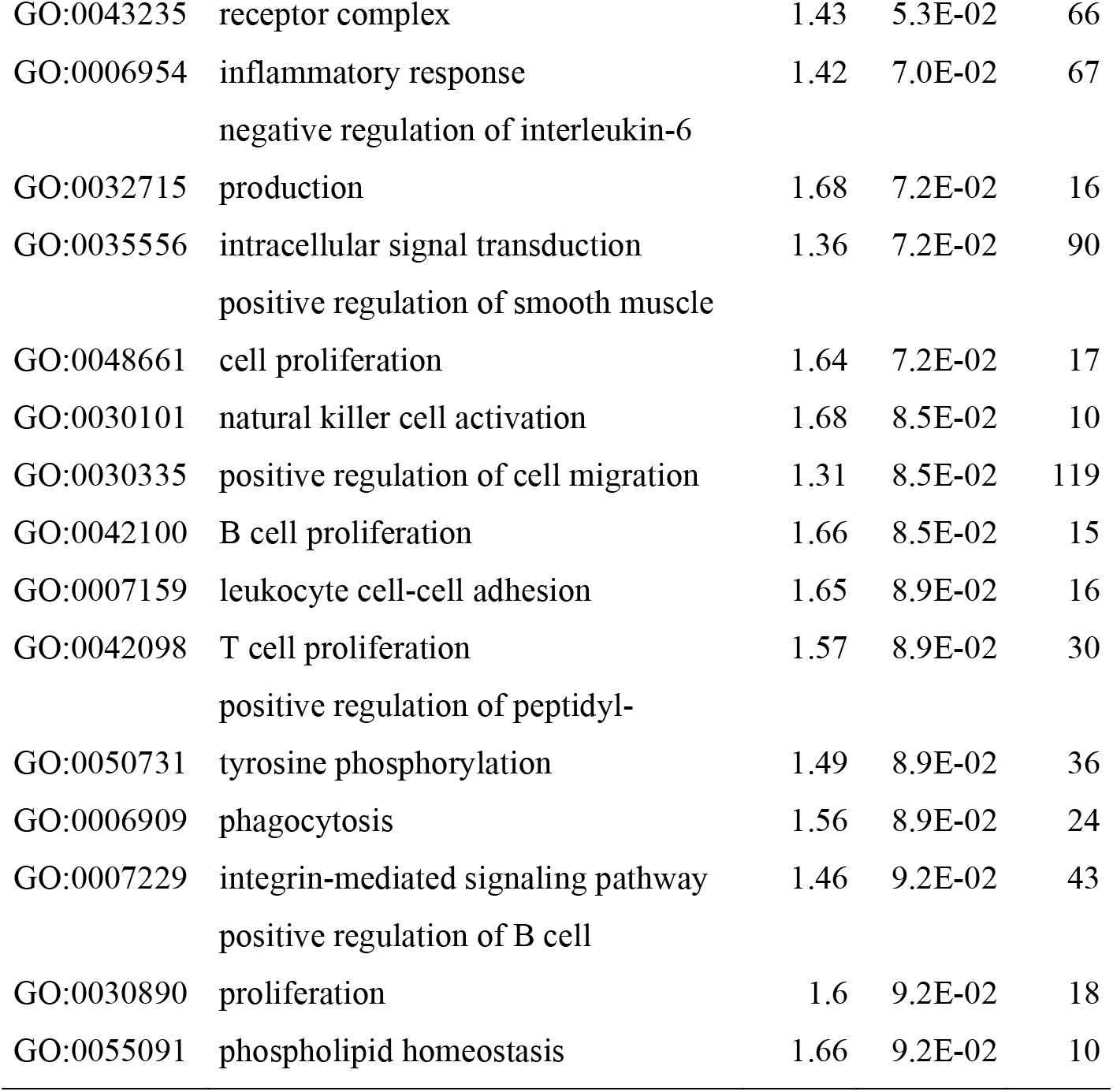
Gene Ontology terms enriched among genes with low p-values for a model testing association between gene expression and infection intensity.

**Figure 2.**
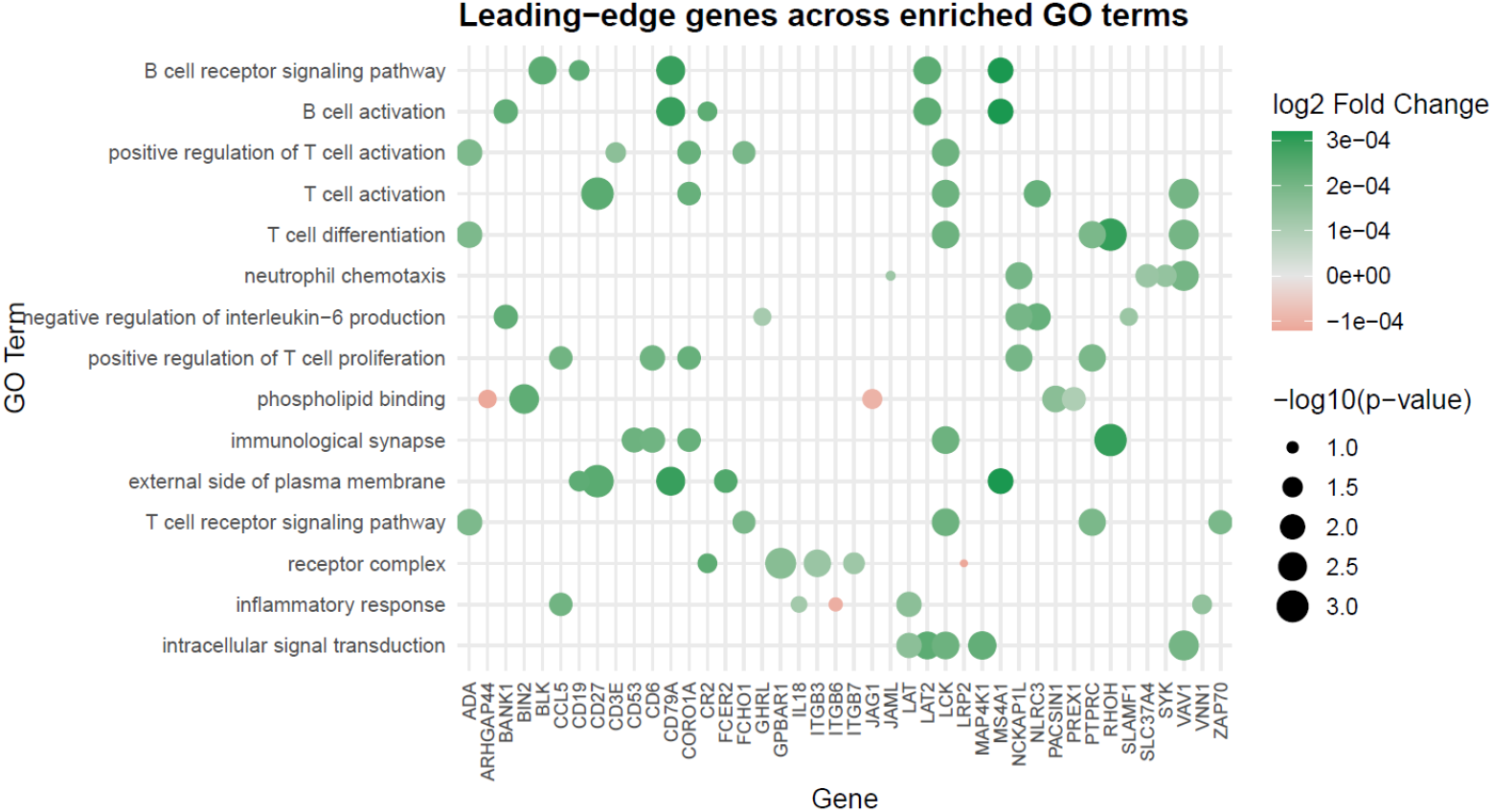
Leading-edge genes across enriched GO terms. For each category top five genes are selected based on the largest absolute values of log2FC. Genes with positive associations between infection intensity and expression are shown in green, and those with negative associations are in red.

## Discussion

Mounting an appropriate response to parasite infection is essential for maintaining host health and fitness. The strength of this response—encompassing immediate immune activation, inflammation, and tissue repair—is expected to depend on multiple factors, including parasite characteristics, environmental conditions, and host-specific traits. In this study, we found that the immune response of European bison correlates with the intensity of infection by *A. sidemi*. Although this response is detectable primarily at the level of molecular pathways, the enrichment of these pathways—together with the functional characteristics of the contributing genes—elucidates molecular mechanisms associated with infection.

Typical response to gastrointestinal infection involves stimulation of Th2 immune response to produce their cytokines mediators IL-4, IL-5, IL-9, IL-10 and IL-13 as well as induce IL-5 and IL-13 that eliciting eosinophils and mucosal mast cell effector mechanisms, as well as B-cell recruitment and stimulation to produce antibodies (mainly IgA, IgE) (Hendawy 2018). This pattern has been demonstrated across experimental models as well as human and livestock studies (Grencis 1997; Balic et al. 2000; Maizels et al. 2012). Despite close evolutionary relationships among main gastrointestinal nematode of ruminants, there are, however, significant differences in the parasitic life cycles which impact on the immunity or immunopathology induced by each species. These differences are mainly related to the environmental niches within the host occupied by the larval stages and the mode of feeding of the adult nematodes (Balic et al. 2000). For example, larval stages of mucosal grazers like *Ostertagia spp*. and *Teladorsagia spp*. develop in modified gastric pits which exclude eosinophils, while blood-sucking *Haemonchus contortus* induces eosinophil-mediated attack and damage (Rainbird et al. 1998; Balic et al. 2000). In the case another blood-feeder studied by us here, *A. sidemi*, expression of typical antihelminth interleukins was generally low and we did not detect clear canonical Th2 polarisation. We did, however, detect enrichment of B cell receptor signalling pathway, activation and differentiation, consistent with their role in combating gut parasites in ruminants (Tuo et al. 2016). T-cell signalling pathway, activation and differentiation signals were also detected, with CD4 expression indicating their helper profile. Their involvement in orchestrating immune response, such as leukocyte recruitment and activation and tissue repair is indicated by, respectively, increased expression of VAV1/VAV3, CD3E (Tybulewicz 2005) and TGFB (Deng et al. 2024).

Furthermore, we detected increased expression of genes involved in inflammatory response and phagocytosis. For example, we recorded increased expression of a pro-inflammatory chemokine CCL5 (RANTES) that recruits immune cells, including eosinophils, to sites of inflammation (E McGovern and H Wilson 2013). We also noted activation of tissue-damage response pathway (upregulation of increased expression of MYD88 and HMGB1 (Piccinini and Midwood 2010), as well as increased expression of TGFB, a key cytokine stimulating tissue repair (Deng et al. 2024). These results are consistent with tissue damage reported by Osińska and Demiaszkiewicz (2010), demonstrating pathological lesions in individuals infested with *A. sidemi* as infiltrations of inflammatory cells in the walls of abomasa and duodena (mainly lymphoid cells and eosinophils), hyperaemia, oedema, mucosal lesions and proliferations of lymphatic follicles at various levels of intensity We detected a correlation between infection level and expression of several genes involved in negative regulation of interleukine-6, a multifunctional cytokine involved in the inflammatory response production. This correlation may reflect host modulation of immune response to limit tissue damage (Estrada-Reyes et al. 2015).

Overall, our results suggest that the intensity of the immune response increases with infection level, consistent with expectations for parasite-driven activation. Although we detected several pathways associated with anti-helminth immunity, the canonical transcriptional signature of Th2 polarisation was notably absent, raising the possibility that an impaired Th2 response may limit effective parasite control. This interpretation is consistent with experimental studies, where deficiencies in Th2-associated interleukins lead to substantially higher worm burdens during both acute and chronic infections (Volkmann et al. 2003; Licona-Limón et al. 2013). However, despite the lack of overt Th2 cytokine induction, we observed relatively high expression of IL4R and STAT6 (logCPM = 4.3 and 5.3, respectively)—core components required for Th2 differentiation (Finkelman et al. 2004). This suggests that the abomasal mucosa likely remains competent to respond to type-2 cytokine signaling, even if the signaling cascade is not strongly transcriptionally activated under the conditions measured.

A potential fitness consequence of these immunological effects of infection with *A. sidemi* is that they may drive evolutionary change in the host immunome. Such change, however, is possible only when functional genetic variation exists in the underlying immune genes. The European bison population underwent severe historical bottlenecks and consequently exhibit low overall genetic diversity. Genome-wide analyses remain, however, limited, and detailed studies of immune-related loci have focused only on a small set of candidate genes (Radwan et al. 2010; Babik et al. 2012). Our results therefore provide an essential foundation for future work aimed at characterizing variation in immune genes that may be particularly relevant in the context of emerging parasites.

## Acknowledgements

We thank Barbara Marczuk, Ewelina Hapunik and Dariusz Chilecki from MRI PAS for collection and preserving European bison tissue samples and Sylwia Jedut for help with RNA extractions.

## Funding

This work was supported by the “Initiative of Excellence - Research University” AMU grant to MK (140/04/POB1/0001). The material was collected in the frame of the project no. 2012/07/B/NZ8/00066 financed by the National Science Centre.

## Author Contributions

All authors contributed to the study conception and design. Sampling, material preparation and data collection was performed by M. K.-S. and R.K. Data analyses were performed by M.K. and results were interpreted by M.K. and J.R. The first draft of the manuscript was written by M.K. and J.R. and all authors commented on the previous version of the manuscript. All authors read and approved the final version of the manuscript.

## Competing interests

The authors declare no competing interests.

## Data availability

Raw sequence data have been deposited in the NCBI Sequence Read Archive (BioProject PRJNA1373505) and will be available upon publication.

## Notes

### Competing Interest Statement

The authors have declared no competing interest.

